# Discovering the shared biology of cognitive traits determined by genetic overlap

**DOI:** 10.1101/723643

**Authors:** JPOFT Guimaraes, J Bralten, CU Greven, B Franke, E Sprooten, CF Beckmann

## Abstract

Investigating the contribution of biology to human cognition has assumed a bottom-up causal cascade where genes influence brain systems that activate, communicate, and ultimately drive behavior. Yet few studies have directly tested whether cognitive traits with overlapping genetic underpinnings also rely on overlapping brain systems. Here, we report a step-wise exploratory analysis of genetic and functional imaging overlaps among cognitive traits. We used twin-based genetic analyses in the human connectome project (HCP) dataset (N=486), in which we quantified the heritability of measures of cognitive functions, and tested whether they were driven by common genetic factors using pairwise genetic correlations. Subsequently, we derived activation maps associated with cognitive tasks via functional imaging meta-analysis in BrainMap (N=4484), and tested whether cognitive traits that shared genetic variation also exhibited overlapping brain activation. Our genetic analysis determined that six cognitive measures (card sorting, no-go continuous performance, fluid intelligence, processing speed, reading decoding and vocabulary comprehension) were heritable (0.3<h^2^<0.5), and genetically correlated with at least one other heritable cognitive measure (0.2<ρ_g_<0.35). The meta-analysis showed that two genetically-correlated traits, card sorting and fluid intelligence (ρ_g_=0.24), also had a significant brain activation overlap (*ρ*_perm_=0.29). These findings indicate that fluid intelligence and executive functioning rely on overlapping biological features, both at the neural systems level and at the molecular level. The cross-disciplinary approach we introduce provides a concrete framework for data-driven quantification of biological convergence between genetics, brain function, and behavior in health and disease.

## Introduction

The current biological understanding of cognition relies on an assumption of a bottom-up cascade of genetic variants, that affect cell functions, brain activation and connectivity, and ultimately converge onto behavior (1–4). As big data and meta-analytic resources become increasingly available, new opportunities arise to understand how different domains of human biology converge and lead to interindividual differences in cognitive ability. Still, few studies to date have directly investigated the correspondence across three levels of biological complexity, integrating cognitive constructs, brain function, and their genetic underpinnings (Figure 1).

**Figure 1.**
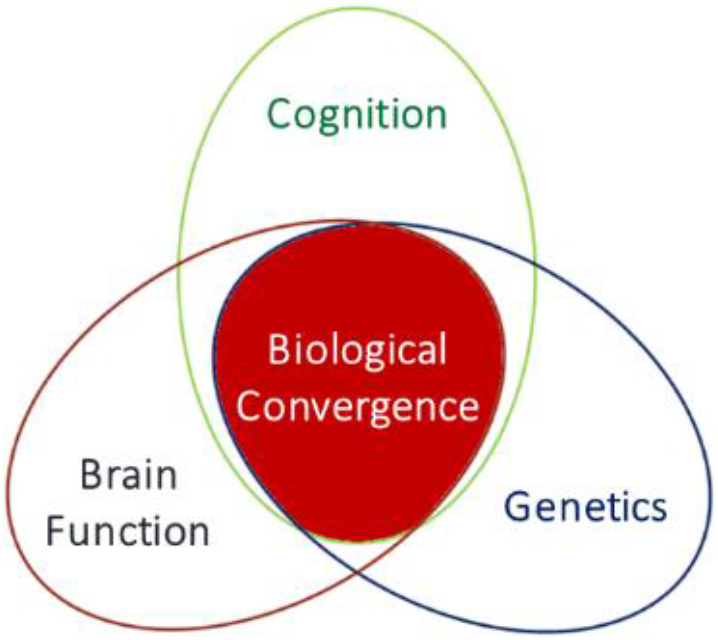
Venn diagram picturing the biological convergence across the domains of cognition, brain function, and genetics. This convergence represents the biological factors involved in the bottom-up cascade that crosses multiple levels of human biology, and ultimately have influence on cognitive performance. In this study, we report a step-wise and data-driven approach that aims to investigate whether this biological convergence may be shown across many traits of cognitive performance, by looking at their overlapping genetic factors and brain activation patterns.

Cognition is an umbrella term for a range of higher-order functions, whose genetic underpinnings have been investigated by means of family studies in general, and twin studies in particular (5, 6). Structural equation modeling (SEM) of twin-pair relationships (i.e. twin modeling) has consistently shown that cognitive measures, such as verbal and non-verbal intelligence, perceptual speed, working memory, reading, and math skills are significantly heritable, i.e. driven by additive and non-additive genetic variation (5, 7–9). Furthermore, twin studies have shown that cognitive measures correlate genetically, that is they share some of their genetic underpinnings, and that a proportion of their phenotypic correlations is significantly mediated by shared genetic variation. These shared genetic underpinnings across specific cognitive traits intuitively suggest that these traits rely on overlapping neurobiological systems (i.e. “circuits”).

Using functional magnetic resonance imaging (fMRI), cognitive tasks are routinely found associated with the activation of specific brain regions, generally organized within networks (10–12). Regions within and across these networks are consistently found to be co-activated during the execution of specific tasks, as well as during rest (13, 14). Functional MRI activation patterns during performance of different cognitive tasks have been found to overlap to some degree (15–20), but systematic quantification of this is limited to date. Activation and connectivity of brain networks were shown to be heritable (21–23), and, more recently, activation was found to be genetically correlated with cognitive measures (24). However, the degree to which cognition and brain function are genetically dependent on each other is unknown. Concretely, given a multi-level biological ontology of cognitive constructs and domains, one would expect that cognitive processes with shared genetic determinants also show a high degree of correspondence, both in their fMRI activation and co-activation patterns (11).

In this study, we investigated cognition in terms of genetic, circuits, and behavioral performance levels, with the goal of finding evidence for biological convergence across these biological levels. In the comprehensively phenotyped Human Connectome Project (HCP) cohort (http://humanconnectome.org), firstly, we used twin-based univariate genetic analysis to estimate the heritability of different measures of human cognition available for HCP participants (demographics displayed in Table 1). For heritable constructs we subsequently tested in a bivariate manner, sharing of genetic variation as quantified by genetic correlations. We then performed functional imaging meta-analysis in order to estimate probabilistic maps of brain activation associated with each cognitive trait using BrainMap (25), and tested whether there was spatial overlap in cognitive task-associated circuits of traits with shared genetic factors. Finally, we combined the significances estimated for the genetic and circuit overlaps, and inferred via fixed-effects meta-analysis whether each pair of cognitive traits biologically converged, i.e. formed a meta-overlapped phenotype (MOP). With that, we demonstrate that current publicly-available resources are suitable for addressing biological convergence relevant for cognition. A general description of our methods is represented in Figure 2; more details of our procedure can be found in *Methods & Materials*.

**Table 1.**
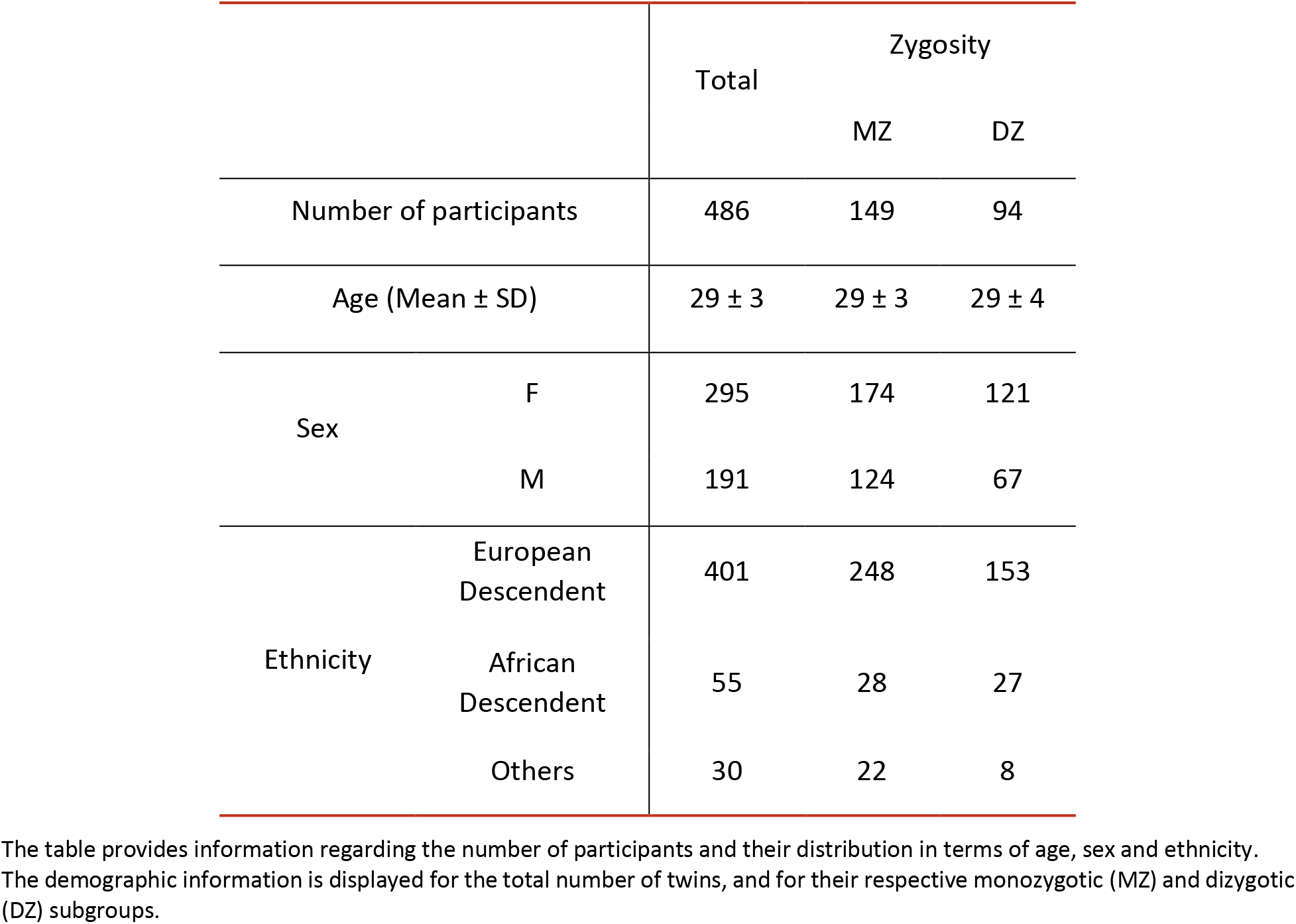
Demographics of HCP participants included in the twin-based genetic analysis.

**Figure 2.**
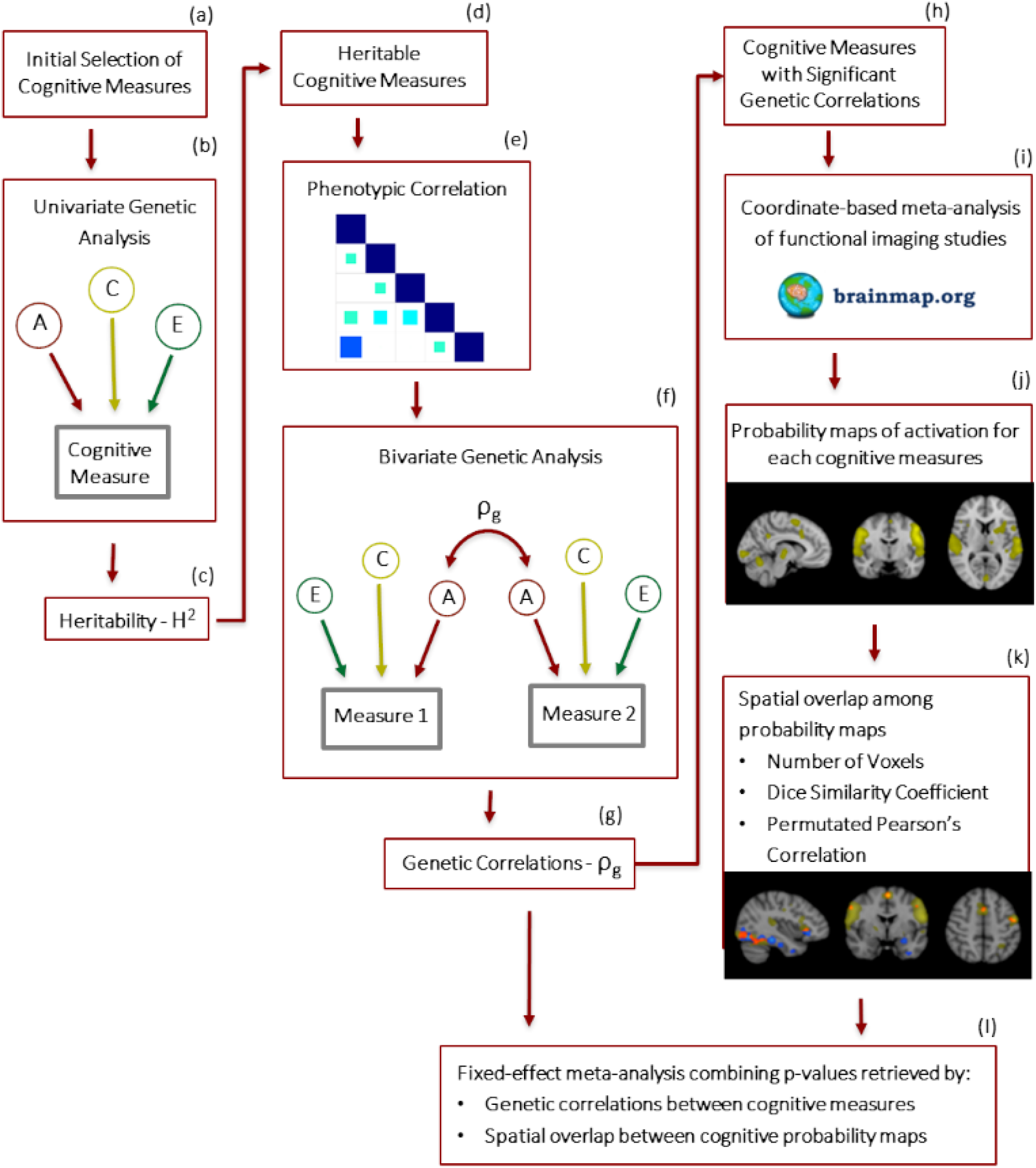
Flow chart summarizing the analysis here implemented. Starting with a selection of cognitive measures evaluating performance among specific higher-order functions (a), we used twin-based univariate genetic analysis (b) to test whether they were driven by genetic variation, i.e. heritable (c). Among heritable cognitive measures (d), we addressed their phenotypic correlations (e), and tested further with a twin-based bivariate genetic analysis (f) whether phenotypic correlations were driven by genetic covariation, i.e. genetically correlated (g). Focusing on cognitive measures that were genetically correlated (h), we conducted for each a coordinate-based meta-analysis based on functional imaging studies available in the BrainMap database (i), retrieving probabilistic activation maps representative of task performance related to these cognitive traits (j). Then, we tested whether these cognitive probability maps overlapped for traits with genetic correlations (k). Finally, we combined p-values retrieved by inferences of genetic correlation and spatial overlap between cognitive maps, and inferred via fixed-effects meta-analysis whether pairs cognitive traits biologically converged based on their overlapping genetics, circuitry, and behavior (l).

## Results

### Univariate genetic analyses

With the data from HCP twin participants (demographics summarized in Table 1), we estimated the heritability of cognitive measures available in the dataset. Heritability estimates (h^2^), and associated statistics of our twin-based univariate analysis, are shown in Table 2. Heritability was significant in the cases of card sorting, no-go continuous performance, delay discounting for 200$ reward, inhibitory control and attention, fluid intelligence, picture sequence memory, processing speed, reading decoding and vocabulary comprehension (p ≤ 0.05). Heritability estimates were moderate in all cases, ranging from 0.3 to 0.5. Additionally, twin correlations of card sorting, no-go continuous performance, inhibitory control and attention, and processing speed indicated the presence of non-additive genetic effects (e.g. dizygotic twin correlation is less than half monozygotic twin correlation), which thus show that these cognitive traits have a broad-sense heritability representative of additive and non-additive genetic effects.

**Table 2.** Univariate genetic results obtained for the fifteen cognitive measures available in the HCP dataset.

| Cognition Measure |  | rMZ (95% CI) | rDZ (95% CI) | h<sup>2</sup> (95% CI) | P(h<sup>2</sup>) | c<sup>2</sup> (95% CI) | e<sup>2</sup> (95% CI) |
| --- | --- | --- | --- | --- | --- | --- | --- |
| Card Sorting |  | 0.4 (0.26-0.52) | 0.1 (-0.11-0.3) | **0.38 (0.07-0.5)** | **0.0213** | 0 (0-0) | **0.62 (0.5-0.75)** |
| Continuous Performance | Go | 0.18 (0.02-0.32) | -0.02 (-0.24-0.19) | 0.15 (0-0.29) | 0.2628 | 0 (0-0.07) | **0.85 (0.71-0.99)** |
|  | No-Go | 0.5 (0.37-0.61) | 0.17 (-0.01-0.34) | **0.49 (0.21-0.6)** | **0.0025** | 0 (0-0.02) | **0.51 (0.4-0.65)** |
| Delay Discounting | 200\$ | 0.48 (0.35-0.59) | 0.26 (0.07-0.42) | **0.45 (0.04-0.59)** | **0.0307** | 0.03 (0-0.38) | **0.52 (0.41-0.65)** |
| | 40k\$ | 0.48 (0.35-0.58) | 0.39 (0.2-0.54) | 0.18 (0-0.56) | 0.3679 | 0.3 (0-0.53) | **0.52 (0.42-0.65)** |
| Fluid Intelligence |  | 0.51 (0.39-0.62) | 0.32 (0.14-0.47) | **0.39 (0.01-0.62)** | **0.0435** | 0.12 (0-0.44) | **0.49 (0.38-0.61)** |
| Inhibitory Control and Attention |  | 0.4 (0.26-0.52) | -0.01 (-0.21-0.2) | **0.36 (0.14-0.49)** | **0.0067** | 0 (0-0) | **0.64 (0.51-0.78)** |
| Picture Sequence Memory |  | 0.49 (0.36-0.6) | 0.24 (0.05-0.4) | **0.49 (0.12-0.6)** | **0.0144** | 0 (0-0.13) | **0.51 (0.4-0.63)** |
| Processing Speed |  | 0.37 (0.23-0.5) | -0.01 (-0.21-0.19) | **0.33 (0.1-0.46)** | **0.0123** | 0 (0-0) | **0.67 (0.54-0.81)** |
| Reading Decoding |  | 0.73 (0.65-0.8) | 0.51 (0.36-0.63) | **0.45 (0.19-0.76)** | **0.0005** | 0.28 (0-0.52) | **0.27 (0.2-0.35)** |
| Spatial Orientation | Correct Position | 0.42 (0.27-0.55) | 0.4 (0.24-0.53) | 0.05 (0-0.45) | 0.797 | **0.37 (0.03-0.51)** | **0.58 (0.45-0.7)** |
|  | Wrong Position | 0.44 (0.29-0.56) | 0.39 (0.23-0.53) | 0.1 (0-0.5) | 0.6211 | 0.34 (0-0.51) | **0.56 (0.44-0.69)** |
| Verbal Episodic Memory |  | 0.35 (0.2-0.49) | 0.25 (0.07-0.41) | 0.2 (0-0.49) | 0.3673 | 0.15 (0-0.41) | **0.65 (0.51-0.8)** |
| Vocabulary Comprehension |  | 0.71 (0.62-0.77) | 0.53 (0.38-0.65) | **0.35 (0.08-0.67)** | **0.0088** | **0.36 (0.05-0.59)** | **0.29 (0.23-0.38)** |
| Working Memory |  | 0.56 (0.44-0.66) | 0.43 (0.26-0.57) | 0.26 (0-0.62) | 0.1416 | 0.3 (0-0.57) | **0.44 (0.34-0.56)** |

The following estimates are shown for each trait: MZ and DZ twin correlation of the trait (rMZ and rDZ, respectively); proportion of variance in the trait explained by genetic effects (i.e. heritability), and their respective 95% confidence interval (CI) estimations (h<sup>2</sup>); significance of the genetic component of the twin modeling (P(h<sup>2</sup>)); proportion of variance in the trait due to shared/common and unique environment, together with their 95% CI estimations (c<sup>2</sup> and e<sup>2</sup>, respectively).
We highlight in **bold** the h<sup>2</sup>, c<sup>2</sup> and e<sup>2</sup> components of variance that were significant in each cognitive measure, inferred based on the p-value of h<sup>2</sup>, and the CI of c<sup>2</sup> and e<sup>2</sup>.

In Table S1, we show that genetic results obtained only with the ethnic majority group (European-descendent subjects, who represent 83% of the total sample) were consistent with the results observed for the whole sample. Heritability of all the cognitive measures remained significant or close to significant (p < 0.2). The lower statistical power associated with modeling a subset of the total sample may explain the lack of significance for some of the cognitive measures.

### Bivariate genetic analyses

Twenty-nine out of 36 phenotypic correlations investigated among heritable cognitive measures were significant (p ≤ 0.05), whose coefficients are shown in Figure S1. Of the 29 pairs of cognitive traits, 25 were successfully modeled in our twin-based bivariate analysis, which reported, without exception, positive phenotypic and genetic correlations for all the pairs (Table 3). The phenotypic correlation coefficients ranged between 0.08 and 0.67. Furthermore, five of these genetic correlations were found to be significant as the 95% confidence interval (CI), in which the lower boundary of the CI did not cross zero: card sorting with fluid intelligence and processing speed, and reading decoding with fluid intelligence, no-go continuous performance, and vocabulary comprehension.

**Table 3.** Correlation matrix showing the bivariate genetic results obtained for the nine heritable cognitive traits, with a priori certified significant phenotypic correlations (p ≤ 0.05).

| | $\rho_{\text{phen}}$ (95% CI) | | | | | | | | | |
| --- | --- | --- | --- | --- | --- | --- | --- | --- | --- | --- |
|  |  | Card Sorting | Continuous Performance No-go | Delay Discounting | Fluid Intelligence | Inhibitory Control and Attention | Picture Sequence Memory | Processing Speed | Reading Decoding | Vocabulary Comprehension |
| $\rho_g$<br>(95% CI) | Card Sorting | | – | – | **0.18**<br>(0.09 - 0.27) | – | **0.21**<br>(0.12 - 0.3) | **0.4**<br>(0.32 - 0.48) | **0.26**<br>(0.17 - 0.35) | **0.12**<br>(0.02 - 0.21) |
|  | Continuous Performance No-go | – |  | – | **0.28**<br>(0.19 - 0.37) | – | **0.11**<br>(0.01 - 0.21) | – | **0.31**<br>(0.22 - 0.4) | **0.2**<br>(0.1 - 0.29) |
|  | Delay Discounting | – | – |  | **0.22**<br>(0.13 - 0.31) | – | – | – | – | **0.27**<br>(0.17 - 0.36) |
|  | Fluid Intelligence | **0.24**<br>(0.01 - 0.38) | 0.12<br>(-0.12 - 0.31) | 0.05<br>(-0.18 - 0.3) |  | – | **0.36**<br>(0.27 - 0.44) | – | **0.45**<br>(0.37 - 0.53) | **0.42**<br>(0.33 - 0.5) |
|  | Inhibitory Control and Attention | – | – | – | – |  | **0.21**<br>(0.12 - 0.3) | **0.3**<br>(0.21 - 0.38) | **0.16**<br>(0.07 - 0.25) | **0.19**<br>(0.09 - 0.28) |
|  | Picture Sequence Memory | 0.2<br>(-0.03 - 0.37) | 0.09<br>(-0.13 - 0.26) | – | 0.11<br>(-0.12 - 0.36) | 0.15<br>(-0.04 - 0.3) |  | **0.18**<br>(0.08 - 0.27) | **0.22**<br>(0.13 - 0.32) | **0.21**<br>(0.11 - 0.31) |
|  | Processing Speed | 0.25<br>(0.06 - 0.37) | – | – | – | 0.11<br>(-0.02 - 0.28) | 0.18<br>(-0.01 - 0.33) |  | **0.19**<br>(0.1 - 0.28) | **0.15**<br>(0.06 - 0.25) |
|  | Reading Decoding | 0.19<br>(-0.04 - 0.39) | **0.27**<br>(0.08 - 0.45) | – | **0.26**<br>(0.05 - 0.51) | 0.06<br>(-0.15 - 0.25) | -0.01<br>(-0.2 - 0.22) | 0.09<br>(-0.11 - 0.29) |  | **0.67**<br>(0.61 - 0.72) |
|  | Vocabulary Comprehension | 0.1<br>(-0.13 - 0.32) | 0<br>(-0.19 - 0.21) | 0.03<br>(-0.17 - 0.27) | 0.12<br>(-0.07 - 0.35) | 0.14<br>(-0.07 - 0.33) | 0.03<br>(-0.19 - 0.28) | 0.17<br>(-0.05 - 0.35) | **0.32**<br>(0.12 - 0.57) |  |

In the entries positioned within the upper right triangle, we see the phenotypic correlations estimated by the twin model ( $\rho_{\text{phen}}$ ), and their respective 95% confidence interval (CI) estimates.
Within the bottom left triangle, the entries show the estimates for genetic correlations between cognitive traits ( $\rho_g$ ), and the respective 95% CI inference, in which we highlight **in bold** the significant genetic correlations (e.g. the estimates for which the lower boundary of the CI did not cross zero).
Combinations of traits for which phenotypic correlations prior twin modeling were not significant are signaled with a dash ("–"), so as the genetic correlations that could not be modeled.

### Functional imaging meta-analysis

We derived individual probabilistic maps of activation for five of the individual cognitive traits that we found to be genetically correlated (see Table 3). We could not perform a meta-analysis of processing speed due to the lack of functional imaging studies in the BrainMap database reporting an HCP-equivalent task performance. Main information regarding the meta-analysis conducted for each trait are presented in Table 4, whereas additional details are reported in Table S2. Moreover, the studies and respective activation contrasts collected for the meta-analyses of card sorting, no-go continuous performance, fluid intelligence, reading decoding, and vocabulary comprehension are listed separately for each cognitive trait in Tables S3-S7.

**Table 4.** Information regarding the meta-analytic search conducted for cognitive traits individually found to be involved significant genetic correlations.

| Cognition measure | Number of Studies | Number of Samples | Total number of subjects (number of healthy subjects) |
| --- | --- | --- | --- |
| Card Sorting | 26 | 33 | 678 (591) |
| Continuous Performance<br>No-go | 64 | 98 | 1705 (1255) |
| Fluid Intelligence | 18 | 29 | 417 (372) |
| Reading Decoding | 28 | 33 | 385 (343) |
| Vocabulary Comprehension | 78 | 97 | 1299 (1178) |

From the total number of participants, we highlight for each trait the number of healthy subjects, from whom we collected the majority of the reported coordinates we used in further meta-analytic steps.

### Brain activation overlap

For relevant pairs of cognitive traits with probabilistic activation maps estimated via functional imaging meta-analysis, we evaluated spatial overlap visually (Figure 3, Movies S1-S4), and by means of three quantitative measures (Table 5): number of voxels, which provides an absolute measure of volumetric overlap; DSC, a standard relative measure of volumetric overlap; Pearson’s correlations with permutation-based inference, an inference-based relative measure of surface overlap. The three measures showed consistent results showing a high degree of spatial overlap between card sorting and fluid intelligence, which was the only significant result according to the inference-based approach.

**Table 5.**
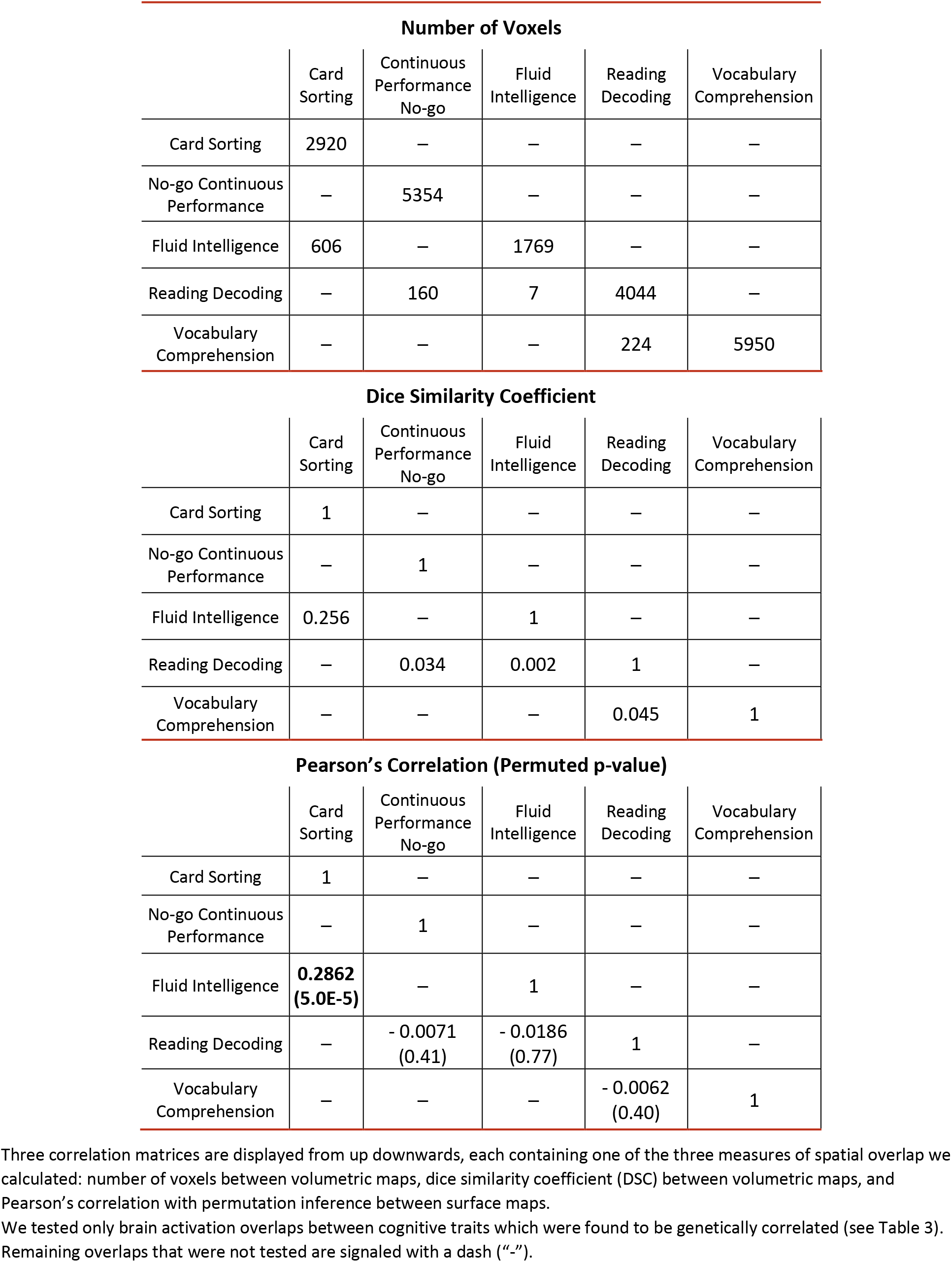
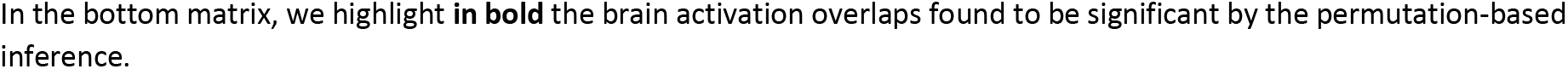
Correlation matrices showing results relative to spatial overlaps between probabilistic activation maps estimated for card sorting, no-go continuous performance, fluid intelligence, reading decoding, and vocabulary comprehension.

**Figure 3.**
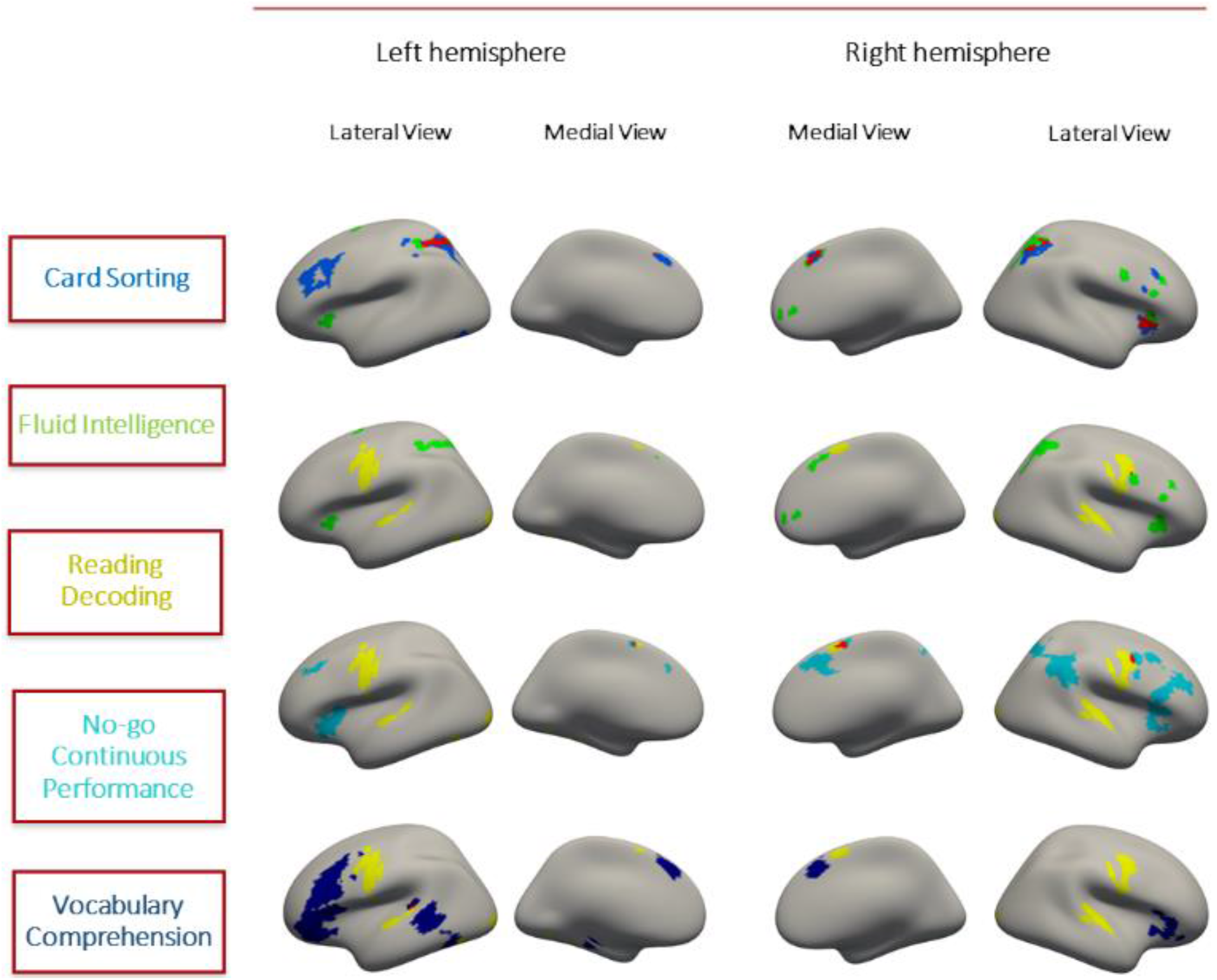
Brain surface visualization of probabilistic activation maps estimated for card sorting, fluid intelligence, reading decoding, no-go continuous performance and vocabulary comprehension (colored in blue, green, yellow, light-blue and dark-blue, respectively). Among these maps, four brain activation overlaps were addressed in result our genetic correlation results (in Table 3), here sorted from top to bottom: card sorting with fluid intelligence, and reading decoding with fluid intelligence, no-go continuous performance and vocabulary comprehension (overlapping regions highlighted in red). Individual and overlapped activations are observed in frontal, parietal and temporal brain regions.

### Biological convergence between cognitive traits

We highlight in Table 6, for each pair of cognitive traits, the p-values retrieved by their genetic and circuit overlap inferences, respectively, and the results of their fixed-effects meta-analysis. Given we tested four pairs of cognitive traits in terms of their overlapping genetics and circuitry, the fixed-effects meta-analysis provided evidence for card sorting and fluid intelligence to be a significant MOP, i.e. to show significant biological convergence in terms of their genetics, circuitry, and behavior (p ≤ 0.01).

**Table 6.** Fixed-effects meta-analysis conducted on p-values retrieved by genetic correlation (P(ρ_g_)) and brain activation overlap P(ρ_perm_) estimated for pairs of cognitive traits that were tested in terms overlapping genetics and circuitry.

| Pairs of Cognitive Traits | | $P(\rho_g)$ | $P(\rho_{perm})$ | $P(\text{Fisher's})$ |
| --- | --- | --- | --- | --- |
| Card Sorting | Fluid Intelligence | **0.033** | **4.8e-5** | **2.4e-5** |
| Reading Decoding | Fluid Intelligence | **0.063** | 0.77 | 0.20 |
| Reading Decoding | No-go Continuous Performance | **0.016** | 0.41 | 0.040 |
| Reading Decoding | Vocabulary Comprehension | **0.019** | 0.40 | 0.045 |

We tested the following four pairs of cognitive traits: card sorting paired with fluid intelligence, and reading decoding with fluid intelligence, no-go continuous performance, and vocabulary comprehension.
We display the significances of overlap-infering test, and the results of the respective fixed effect meta-analysis that was conducted on the two previously referred p-values, for each cognitive trait pair.
Fixed-effect meta-analysis was implemented using Fisher's sum of logs method, which provides a p-value ( $P(\text{Fisher's})$ ) indicating whether each trait pair biologically converged in terms of their genetics and circuitry ( $p \leq 0.01$ ).
Significant p-values are highlighted in **bold** for $P(\rho_g)$ and $P(\rho_{perm})$ ( $p \leq 0.05$ ), and $P(\text{Fisher's})$ ( $p \leq 0.01$ ).

## Discussion

In this study, we investigated the biological convergence across genetics, circuits, and cognition, by means of twin data and meta-analytic resources that are publicly available. We report a significant MOP involving card sorting and fluid intelligence, in which the two cognitive traits biologically converge in terms of their genetics, brain activation, and behavioral performance. Firstly, we showed using twin-based univariate genetic analysis that genetic effects explained performance in nine cognitive measures present in our dataset. Next, in a bivariate genetic analysis, we showed that six cognitive measures shared genetic factors that significantly explained a proportion of five of their phenotypic correlations. Then, we took into account the six genetically-correlated cognitive traits in a functional imaging meta-analysis with BrainMap. We conducted a coordinate-based meta-analysis for each cognitive trait, which successfully retrieved individual probabilistic maps of brain activation for five of the six genetically-correlated traits. Further, we found that card sorting and fluid intelligence were the only pair of cognitive traits with shared genetics that also exhibited a significant spatial overlap between their activation maps. We confirmed afterwards, via fixed-effects meta-analysis of p-values, that this cognitive trait pair was the only to biologically converge.

Our findings for the heritability of the nine cognitive traits are consistent with previous literature (7, 9, 26–35). Our results on genetic correlations between card sorting and fluid intelligence, card sorting and processing speed, and those of reading decoding with the no-go continuous performance, fluid intelligence, and vocabulary comprehension extend existing literature. Previous twin studies have shown that cognitive flexibility, as measured by the card sorting test, shares genetic factors with general cognitive ability (9), and with other, more specific, measures of cognition such as working memory. By taking a more data-driven approach, we showed that card sorting is also genetically correlated with fluid intelligence and processing speed. Regarding the genetic correlations involving reading decoding, previous literature supports a genetic link with general markers of intelligence (7, 31, 36), as well as specific aspects of intelligence, such as problem solving and reading comprehension (26). Therefore, the fact that we found reading decoding to be genetically correlated with fluid intelligence and vocabulary comprehension strengthens the idea of a common genetic background between reading and general intelligence, as well as with the more specific cognitive domain of language. To our knowledge, however, no previous studies have reported a genetic link of reading decoding with continuous performance. Still, this genetic correlation may relate to the idea that reading skills require sustained attention and to its shared genetics with inhibition problems (37, 38), which are commonly captured using go/no-go experimental designs (39).

With our functional imaging meta-analysis, we derived probabilistic maps of activation for cognitive traits showing genetic overlap. By accessing the BrainMap database (25), one of the biggest available to conduct coordinate-based meta-analysis, we collected a minimally-required number of studies for having a sample representative of each trait, and thus a well-powered meta-analysis (40), except for processing speed. Overall, we observed probabilistic maps that reliably captured activation reported across studies, showing individual and overlapped regions of activation located within the frontal, parietal, and temporal lobes. When testing whether cognitive traits sharing genetic factors also reported spatial overlap in their meta-analytic activation maps, we found card sorting and fluid intelligence to show significant overlap, whereas the overlaps involving reading decoding with fluid intelligence, no-go continuous performance, and vocabulary comprehension were not significant. Without addressing the genetics of cognitive traits, other studies had already reported common regions of activation between some of the cognitive traits tested here, which also belonged to frontal, parietal and temporal areas of the brain (16, 19, 41, 42). However, none of these studies tested the overlapping traits we reported in our analysis. The fact that we implemented a data-driven and step-wise approach through genetics did contribute to the discovery of a spatial overlap between card sorting and fluid intelligence, and further reporting of their biological convergence by means of fixed-effects meta-analysis. This finding emphasizes the relevance of conducting exploratory analysis in the search for the biological convergence across genetics, brain function, and cognition.

By using a data-driven approach based on a genetically informed variable selection, we found that most genetically correlated traits did not show significant brain activation overlap. This does not immediately imply that they do not have a shared neurobiology, but rather that any neurobiology they may share is not generally and consistently reflected in, or captured by, fMRI studies. There are many biological levels intermediate between the genetic and brain mesoscopic level, and the biological convergence of these cognitive traits may be better reflected on a more microscopic scale of human neurobiology. Alternatively, biological convergence may still occur at the mesoscopic level, and be captured using other types of imaging modalities looking at different types of brain measurements, such as structure or electrophysiology-related measures. In any case, our current resources are still limited to fully capture how complex traits, such as cognition, are influenced by this bottom-up effect crossing multiple levels of human biology. Still, our positive result shows that in some cases we can capture this convergence at the level of brain activation.

Regarding the genetic methods employed here, we opted for using twin modeling, a standard approach in the field of psychiatric genetics. Our study shows, consistent with existing literature, that using twin modeling is a powerful and reliable tool to estimate the genetic and environmental factors influencing behavior and cognition (5, 7–9). However, we did not find significant heritability for cognitive traits such as spatial orientation, verbal episodic memory, or working memory, contrary to other twin studies (9, 43, 44). These non-findings may be explained by our moderate sample size, and by the assumptions underlying twin modeling and its classic view of multifactorial traits (45). Twin modeling imposes constraints on the data to be analyzed, such as the need to have a trait measure that follows normal distribution; so as conceptual constraints concerning equal environments and gene-environment interplay. Given the knowledge we currently have regarding gene-environmental correlations (46), and environmental influences on epigenetic factors that may alter gene expression (47), it is questionable whether genetic variance based on twin relationships is driven directly by additive variation in the DNA. Alternative methods modeling single nucleotide polymorphism (SNP) data provide direct link to genomic differences between individuals (48–50); they have already been used successfully in estimating heritability and genetic correlations among cognitive traits (50, 51). Still, we were prompted to use twin modeling instead of SNP modeling, because it provides a rather broad-sense heritability that accounts more comprehensively for the effects of genetic variants, including non-additive genetic effects and SNPs not genotyped/imputed(51, 52). Another advantage of the twin design over SNP modeling is, despite the assumptions, the better characterization of environmental factors by disentangling the effects driven by shared and unique environment.

Strengths of our study are: the deep phenotyping of the HCP sample, with a comprehensive set of cognitive measures in a substantial sample size; and the use of twin modeling and functional imaging meta-analysis in a data-driven and step-wise approach, which allowed investigating unprecedented research questions in the fields of cognition, neuroimaging, and genetics. Nonetheless, there are limitations that may be taken into account for future studies following this kind of approaches. One of them relates to the fact that twin modeling does not provide an optimal way to correct for ancestral differences in multi-ethnic samples. Still, we expected our heritability estimates not to be driven by these differences. Our supplementary analysis shows that modeling only the major ethic group provided results pointing to the same conclusion as the main findings we obtained based on the whole sample (Table S1), which indicates a very unlikely influence of multi-ethnicity in our genetic results. Apart from that, we did not intend to investigate any association between our results and ethnicity. In our view, there are worldwide initiatives, with a wider ethnic representation within and across nations, that are better equipped to answer this kind of questions. We see these initiatives, along with the multi-ethnic sample provided by the HCP consortium, as the first step of many toward a better representation of the population we scientists aim to understand.

Another limitation we encountered with our approach was associated with the lack of systematic measurement for quantifying spatial overlap among brain imaging maps. We chose the three measures that, according to our knowledge, were better design to estimate this quantity: number of voxels, DSC, and the recent method inferring Pearson’s correlations with permutation inference (20). The latter method is up to date, as far as we know, the only one that provides statistical inference of spatial overlaps among meta-driven activation maps. Validating methods that infer spatial overlap has been dependent on controversial and still on-going questions, such as the definition of a true null hypothesis for this particular test (53, 54). We raise awareness for debating and clarification of these limitations, and we see the permuted-correlation method we used as a significant progress toward the solution (20).

In conclusion, we find, by means of our data-driven and step-wise approach, evidence pointing to the biological convergence among genetics, brain function, and behavioral performance for selected measures of cognition. This finding points to the idea that cognition results of a bottom-up effect covering the genetic and brain functional levels, and that different types of cognition come from the same effect. The idea of biological convergence, we portrait in this study, emphasizes that a better understanding of cognition involves a deeper knowledge of its biological underpinnings and their respective associations. Furthermore, our demonstration of biological convergence has broader implications for the study of human behavior in health and disease, as it provides a concrete framework towards more precise selection and characterization of traits and their associated neural systems and biological pathways.

## Methods & Materials

### HCP sample

We used the S1200 sample released by the WU-Minn HCP consortium in March 2017 (55). The healthy subjects present in the HCP S1200 sample were recruited from the Missouri Family and Twin Registry. Our sample consisted of 149 monozygotic (MZ) and 93 dizygotic (DZ) twin pairs (from which 92 DZ twin pairs share the same sex), with an age range of 26-32 years old. Twin subjects were included if they had fifteen cognitive scores reflecting task accuracy among the twelve cognitive tasks we used for further analysis: card sorting, go/no-go continuous performance, delay discounting (200$ and 40k$ reward), fluid intelligence, inhibitory control and attention, picture sequence memory, processing speed, reading decoding, spatial orientation (correct and wrong position), working memory, verbal episodic memory, and vocabulary comprehension. Detailed description about these tasks is available in https://wiki.humanconnectome.org/. Demographics about the sample we analyzed is provided in Table 1.

### Twin Modeling

We performed univariate and bivariate genetic analysis of HCP cognitive data by means of twin modeling. We conducted our twin modeling approach using R v3.3.3 (https://www.r-project.org/), and the software package OpenMx v2.8.3 (56).

Prior to modeling, cognitive measures that did not follow a normal distribution, as indicated by a significant Shapiro-Wilk test, were transformed. First, by excluding outliers (+/− 3 standard deviations), and if necessary, via inverse-normal transformation. Where age and sex significantly explained the output variable (determined by linear regression), they were included as covariates in the twin model.

We applied twin models according to the standard ACE design. Based on the genetic and environmental relationships among twins, the ACE model quantifies the extent to which three latent factors drive the variance in a trait: additive genetic (A), shared environmental (C), and unique environmental (E) factors; for a detailed explanation, consult publication by Fruhling and Sham (6). We first ran a univariate ACE model for each cognitive measure to quantify its heritability. The significance of the heritability estimates was estimated by comparing each ACE model with its respective CE model as a null-model (p ≤ 0.05). Next, heritable cognitive measures were further investigated in terms of their phenotypic and genetic correlations. We calculated phenotypic correlations in the whole sample, independent of zygosity, by computing the Pearson’s correlation coefficient and respective significance. For each trait pair with significant phenotypic correlation, we tested with a bivariate ACE model whether the phenotypic correlation was partly driven by shared genetic factors. The significance of the genetic correlation was determined via confidence intervals (CI = 95%).

Additionally, by taking into account that we used a multi-ethnic sample, we intended to show that our genetic results were representative of all the ethnicities here reported. Thus, we designed an extra univariate genetic analysis exclusively with biggest ethnic group (European descents, 83% of the total sample), in order to exclude the hypothesis that our genetic results were driven by ethnicity.

### Functional Imaging Meta-Analysis

For cognitive traits with significant genetic correlations, we performed a coordinate-based meta-analysis to estimate probabilistic activation maps that resembled task performance associated with each of these traits. We conducted our meta-analysis using the database and software tools provided by the BrainMap Initiative (25).

Firstly, we accessed the BrainMap database via Sleuth 2.4 to search for studies whose procedure matched, according to the BrainMap taxonomy (10), the HCP-equivalent cognitive task (last search conducted in September 26, 2018). Each study in the database reported one or more imaging contrasts of conditions (e.g. task versus rest), which yielded the coordinates of interest that we selected for our analysis. In this selection, we accounted for bias related to having different number of contrasts across studies (57). Detailed description of the steps taken in this meta-analytic search is reported in *SI Appendix, Methods: Detailed Description of Meta-Analytic Search.*

Further, based on these coordinates, we computed probabilistic activation maps for each cognitive trait using GingerALE v2.3.6, the activation likelihood estimation (ALE) algorithm provided by BrainMap (40, 58, 59). We used GingerALE to apply cluster-level inference (cluster-forming threshold = 0.001; 10,000 permutations for cluster-size null distribution; cluster-level threshold = 0.05), retrieving thresholded Z-score maps for each cognitive trait. For a detailed description of our ALE settings, consult *SI Appendix, Methods: Settings of Activation Likelihood Estimation Analysis.*

### Brain activation overlap

Given the meta-analytic activation maps we obtained for each cognitive trait, we investigated spatial overlaps between cognitive maps, in order to find whether they had overlapping circuits.

This step in our pipeline was constrained by the current limitations in quantifying and inferring spatial overlaps between volumetric activation maps, with varying smoothness and spatial characteristics, and the lack of established standard methods to perform this type of analysis. An exception to current scenario is the recently reported permutation-based approach using Pearson’s correlation as a measure of spatial overlap (20); method is available online (https://www.github.com/spin-tes). Therefore, we opted for estimating three complementary measures of spatial overlap using Matlab v2018b (https://www.mathworks.com/): (a) the number of imaging voxels in the intersection of the two maps; (b) the Dice similarity coefficient (DSC); and (c) the Pearson’s correlation coefficients with permutation-based inference (20484 permutations).

With the volumetric Z-score maps retrieved by GingerALE, we counted the number of activated voxels in each map to obtain the total number of voxels per cognitive trait, whereas the minimum-statistic map among two cognitive-specific maps were used to estimate the number of overlapping voxels between the traits.

Afterwards, we calculated DSC between maps, given by the mathematical expression:

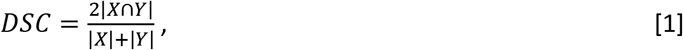

where, |X| and |Y| represent the number of activated voxels in the maps X and Y, respectively, and |X∩Y| the number of activated voxels common to both X and Y. We binarized the Z-maps retrieved by GingerALE (Z-score threshold = 2.3) and, for each cognitive trait pair, we calculated the DSC overlap between their maps (maps X and Y, according to the DSC expression [1]).

For the permuted correlation approach, we projected the thresholded Z-maps retrieved by GingerALE onto the FreeSurfer surface template by nearest neighbor interpolation (fsaverage5, consisted of 10,242 vertices per hemisphere), because this permuted-correlation method does not support volumetric maps. Our cortical surface maps resulted of sampling and averaging across eleven equidistant surface projections in the cortical layer, from the white to the pia mater surfaces. After the volume-to-surface conversion, we built a null distribution by means of 20,484 rotational permutations (two times the number of vertices), and further tested whether the correlation between two surface maps were significant according to the permuted null distribution (p ≤ 0.05).

### Relevance of step-wise pipeline

Our exploratory analysis involved the performance of multiples tests. If all the tests we performed, across all the above steps, were conducted independently from each other, our analysis would be confounded by type I error inflation. Furthermore, not limiting the number of tests would have been particularly unfavorable for estimating genetic correlations, due to the decrease statistical power related to moving from univariate to bivariate genetic analysis. Thus, we used a step-wise approach, in which each analysis step served as a variable selection for the subsequent step. First, only heritable traits were included in the phenotypic correlation analysis. Of those, only trait-pairs that were significantly correlated were tested for genetic correlations. And from this, only traits showing significant genetic correlation, with at least one other trait, were further tested for brain activation overlap.

### Inference of biological convergence

To address whether the pairs of cognitive traits tested throughout our step-wise pipeline represented of MOPs involving biological convergence between cognitive traits, we combined the p-values retrieved by testing genetic correlation and activation overlaps, for each cognitive trait pair, and performed fixed-effects meta-analysis of the two significant values by using the Fisher’s sum of logs method (p ≤ 0.05 / number of cognitive trait pairs). In the end, cognitive trait pairs reporting a significant p-value via this fixed-effects meta-analysis were reported to be biologically converged across the domains of genetics, brain activation and behavior.

## Acknowledgments

J.B. is involved with the PRISM project (www.prism-project.eu<http://www.prism-project.eu>), which has received funding from the Innovative Medicines Initiative 2 Joint Undertaking under grant agreement No 115916. This Joint Undertaking receives support from the European Union’s Horizon 2020 research and innovation programme and EFPIA. This publication reflects only the authors’ views neither IMI JU nor EFPIA nor the European Commission are liable for any use that may be made of the information contained therein. B.F. has received educational speaking fees from Medice and Shire. B.F. received additional funding from a personal Vici grant of the Dutch Organization for Scientific Research (NWO; grant 016-130-669) and from a grant for the Dutch National Science Agenda (NWA) for the NeurolabNL project (grant 400 17 602). E.S. is funded by a NARSAD Young Investigator Award (GRANT ID: 25034), a Hypatia Tenure Track Grant and Christine Mohrmann Fellowship (Radboudumc). Data were provided by the Human Connectome Project, WU-Minn Consortium (Principal Investigators: D.C. Van Essen and K. Ugurbil; 1U54MH091657) funded by the 16 NIH Institutes and Centers that support the NIH Blueprint for Neuroscience Research; and by the McDonnell Center for Systems Neuroscience at Washington University. C.F.B. gratefully acknowledges funding from the Netherlands Organisation for Scientific Research (NWO-Vidi 864.12.003), and the Wellcome Trust UK Strategic Award [098369/Z/12/Z]. C.F.B. gratefully acknowledges support from the Netherlands Organisation for Scientific Research under the Gravitation Programme Language in Interaction (grant 024.001.006).

