## Supplementary Information for "Discovering the shared biology of cognitive traits determined by genetic overlap"

Joao P.O.F.T. Guimaraes, Janita Bralten, Corina U. Greven, Barbara Franke, Emma Sprooten, Christian F. Beckmann

Joao P.O.F.T. Guimaraes


**This PDF file includes:**

Supplementary text
Figs. S1
Tables S1 to S7
Captions for movies S1 to S4

**Other supplementary materials for this manuscript include the following:**

Movies S1 to S4

**Supplementary Information Text**

**Detailed description of meta-analytic search.** We accessed the BrainMap database via Sleuth 2.4 to search for studies whose procedure matched, according to the BrainMap taxonomy, the HCP-equivalent cognitive task (last search conducted in September 26, 2018).

For clarity, here we use “study” to refer to a study described in a single article referenced in BrainMap. We use “group” to refer to a unique sample or subsample within a study (sometimes these are for example controls and patients). And we use “contrast” to refer to the functional MRI (fMRI) or positron emission tomography (PET) contrast of conditions (e.g. task versus rest) of a group within a study that yielded the coordinates provided by BrainMap.

This search consisted of 3 steps: (a) literature search, in which we defined the cognitive terms that best described the HCP-equivalent cognitive task according to the BrainMap taxonomy; (2) literature selection, in which we selected fMRI and PET studies whose experimental design captured an equivalent cognitive performance being measured via the HCP cognitive protocol conducted for a given task; and (3) activation contrast selection, in which we focused on those imaging contrasts that best captured brain activation for a given cognitive trait.

In literature search, We defined a single term relative to the general behavioral domain or the specific task design of the HCP-equivalent cognitive task being searched for each cognitive trait. Table S2 displays the cognitive terms we used to conduct literature search of card sorting, no-go continuous performance, fluid intelligence, reading decoding and vocabulary comprehension.

In the selection of literature and activation contrast, from the studies retrieved by the literature search we conducted for each cognitive trait via the BrainMap database, we selected the imaging contrasts of conditions that yielded the coordinates of interest for our meta-analysis. Each study in the database reported one or more imaging contrasts. We chose one or more activation contrast against a control or baseline condition retrieved by first-level analysis. Table S2 lists the general criteria for selecting activation contrasts representative of card sorting, no-go continuous performance, fluid intelligence, reading decoding and vocabulary comprehension. Furthermore, the list of activation contrasts and respective studies collected individually for each cognitive trait are displayed in the tables S3-S7.

In this selection, we accounted for bias related to having different number of contrasts across studies in this selection. Therefore, it was ensured that every group, and by extension every subject, is contributing an equal amount to the meta-analysis. Furthermore, we took into account that a single study reported activation contrasts for more than one group of subjects, which could differ in terms of the following conditions: health condition, medication, on-going treatment, short-term effects of product ingestion, substance use, IQ level, mother tongue, experimental protocol, age, family relatedness, genotype, training stage, or sex. Under this scenario, we selected activation contrasts reported for one or more group of subjects, independently of their condition. However, we excluded second-level contrasts between subject groups.

**Settings of activation likelihood estimation analysis.** Following the meta-analytic search, we computed probabilistic activation maps for each cognitive trait using GingerALE v2.3.6, the activation likelihood estimation (ALE) algorithm provided by BrainMap. GingerALE computes probability maps reporting the likelihood of brain voxel-level activation across studies given the provided coordinates, and then allows using cluster-wise statistics for whole-brain inference. Throughout this process, we used the additive ALE method in which the meta-analytic network results of computing the union of coordinates reported for each study, and the undilated version of the MNI template provided by the BrainMap database. We estimated the probabilistic maps of brain activation by performing a cluster-level inference with a cluster-forming threshold of 0.001, 10,000 permutations to build the cluster-size null distribution, and a cluster-level threshold of 0.05.

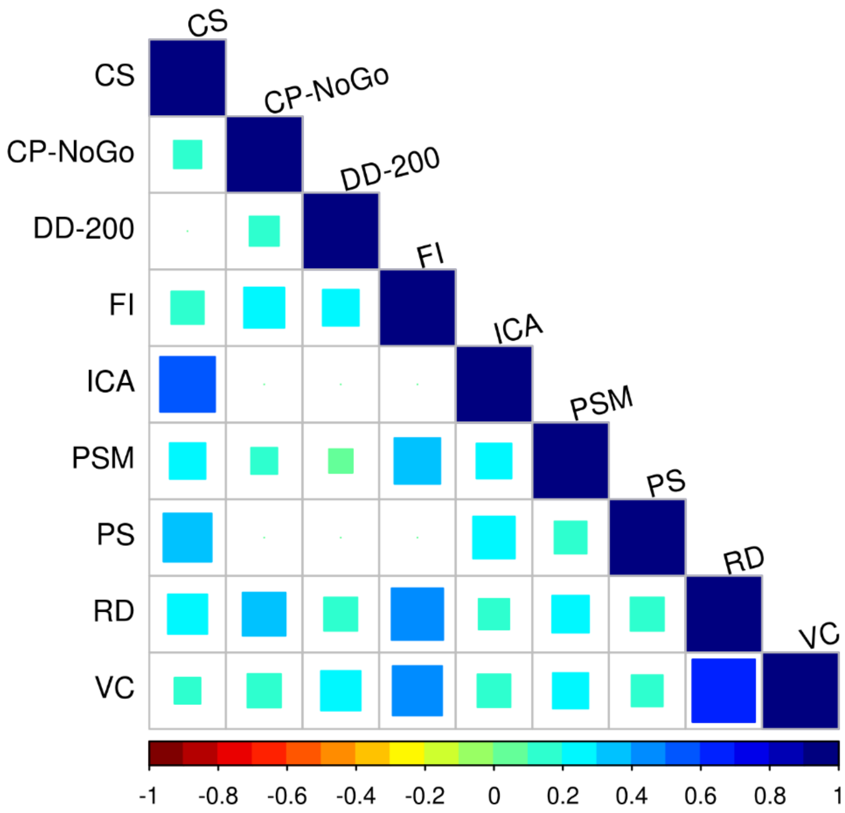


**Fig. S1.** Phenotypic correlations between heritable cognitive traits. Pearson correlation analysis was performed between the following cognitive traits: card sorting (CS), no-go continuous performance (CP-NoGo), delay discounting (DD-200), fluid intelligence (FI), inhibitory control and attention (ICA), picture sequence memory (PSM), processing speed (PS), reading decoding (RD), and vocabulary Comprehension (VC). Correlation coefficients are represented by squares colored based on the color scale coding for the correlation range, in which only significant correlations are plotted.

**Table S1. Univariate genetic results obtained for the fifteen cognitive measures available in the HCP dataset, using a subset consisted only of individuals belonging to the ethnic majority group in the sample (European descendent).**

| Cognition Measure | | rMZ (95% CI) | rDZ (95% CI) | $h^2$ (95% CI) | P($h^2$) | $c^2$ (95% CI) | $e^2$ (95% CI) |
|---|---|---|---|---|---|---|---|
| Card Sorting | | 0.36 (0.21-0.49) | 0.08 (-0.18-0.32) | 0.35 (0-0.48) | 0.0684 | 0 (0-0) | 0.65 (0.52-0.8) |
| Continuous Performance | Go | 0.1 (-0.07-0.27) | -0.02 (-0.25-0.21) | 0.08 (0-0.24) | 0.5734 | 0 (0-0.06) | 0.92 (0.76-1) |
| | No-Go | 0.41 (0.24-0.54) | 0.1 (-0.11-0.3) | **0.38 (0.06-0.52)** | **0.0272** | 0 (0-0.09) | 0.62 (0.48-0.78) |
| Delay Discounting | 200$ | 0.45 (0.3-0.57) | 0.3 (0.08-0.48) | 0.3 (0-0.57) | 0.1931 | 0.15 (0-0.48) | 0.55 (0.43-0.7) |
| | 40k$ | 0.46 (0.32-0.58) | 0.42 (0.22-0.58) | 0.08 (0-0.53) | 0.6945 | 0.38 (0-0.55) | 0.54 (0.42-0.67) |
| Fluid Intelligence | | 0.47 (0.33-0.59) | 0.28 (0.07-0.46) | 0.38 (0-0.59) | 0.0966 | 0.09 (0-0.46) | 0.53 (0.41-0.67) |
| Inhibitory Control and Attention | | 0.39 (0.23-0.52) | -0.04 (-0.27-0.19) | **0.34 (0.09-0.48)** | **0.0166** | 0 (0-0) | 0.66 (0.52-0.81) |
| Picture Sequence Memory | | 0.42 (0.26-0.55) | 0.13 (-0.08-0.32) | **0.4 (0.09-0.53)** | **0.0287** | 0 (0-0.05) | 0.6 (0.47-0.75) |
| Processing Speed | | 0.37 (0.21-0.51) | -0.04 (-0.26-0.19) | **0.33 (0.08-0.47)** | **0.0195** | 0 (0-0) | 0.67 (0.53-0.83) |
| Reading Decoding | | 0.71 (0.62-0.78) | 0.42 (0.22-0.57) | **0.59 (0.26-0.78)** | **3.00E-04** | 0.12 (0-0.43) | 0.29 (0.22-0.38) |
| Spatial Orientation | Correct Position | 0.37 (0.21-0.5) | 0.3 (0.07-0.49) | 0.13 (0-0.5) | 0.6 | 0.23 (0-0.46) | 0.63 (0.5-0.78) |
| | Wrong Position | 0.39 (0.23-0.52) | 0.35 (0.13-0.52) | 0.09 (0-0.51) | 0.7115 | 0.3 (0-0.48) | 0.61 (0.48-0.75) |
| Working Memory | | 0.35 (0.17-0.49) | 0.18 (-0.03-0.36) | 0.34 (0-0.49) | 0.182 | 0.01 (0-0.36) | 0.65 (0.51-0.83) |
| Verbal Episodic Memory | | 0.63 (0.52-0.72) | 0.42 (0.22-0.58) | **0.42 (0.06-0.71)** | **0.0209** | 0.21 (0-0.53) | 0.37 (0.28-0.48) |
| Vocabulary Comprehension | | 0.56 (0.43-0.67) | 0.42 (0.22-0.57) | 0.29 (0-0.65) | 0.1309 | 0.27 (0-0.57) | 0.44 (0.33-0.57) |

The following estimates are shown for each trait: MZ and DZ twin correlation of the trait (rMZ and rDZ, respectively); proportion of variance in the trait explained by additive genetic effects (i.e. heritability), and their respective 95% confidence interval (CI) estimations ($h^2$); significance of the additive genetic component of the twin modeling (P($h^2$)); proportion of variance in the trait due to shared/common and unique environment, together with their 95% CI estimations ($c^2$ and $e^2$, respectively).
We highlight in **bold** the $h^2$, $c^2$ and $e^2$ components of variance that were significant in each cognitive measure, inferred based on the p-value of $h^2$ and the CI of $c^2$ and $e^2$.

**Table S2. BrainMap taxonomy terms used to conduct literature search of card sorting, no-go continuous performance, fluid intelligence, reading decoding and vocabulary comprehension.**

| Cognition Measure | BrainMap Taxonomy Search Term | Criteria for Activation Contrast Selection |
|---|---|---|
| Card Sorting | "Paradigm Class: Wisconsin Card Sorting Test (WCST)" | Decision and switching across dimensions that validate a correct hit |
| No-Go Continuous Performance | "Paradigm Class: Go/No-Go" | Inhibition to no-go signal in a task go/no-go task design |
| Fluid Intelligence | "Paradigm Class: Reasoning/Problem Solving" | Problem solving tasks that typically do not require previous acquisition of verbal knowledge |
| Reading Decoding | "Paradigm Class: Reading (Overt)" | Overt pronouncing of words asked via stimuli |
| Vocabulary Comprehension | "Behavioral Domain is Cognition: Language Semantics" | Semantic categorization and/or matching based on given stimuli |

The cognitive term employed for each cognitive trait is representative of the general behavioral domain (i.e. "Behavioral Domain is Cognition" in the BrainMap taxonomy) or the specific task design of the HCP-equivalent cognitive task (i.e. "Paradigm Class") being searched for each cognitive trait.

**Table S3. List of experiments included for meta-analysis of activation coordinates reported for card sorting task performance.**

| BrainMap ID | Group Type | Group Size | Experiment |
|---|---|---|---|
| 30056 | Healthy Controls | 11 | Conceptual Reasoning - Control |
| 5040008 | Healthy Controls | 7 | Three Dimensional - (Two + One Dimensional) |
| 5040009 | Healthy Controls | 16 | (Negative and Positive Feedback) - (Positive feedback only) |
| 5040011 | Healthy Controls | 11 | Matching After Negative Feedback minus Control Matching (Increases) |
| 5040011 | Healthy Controls | 11 | Matching After Negative Feedback minus Control Matching (Decreases) |
| 5040011 | Healthy Controls | 11 | Matching After Positive Feedback minus Control Matching (Increases) |
| 5040011 | Healthy Controls | 11 | Matching After Positive Feedback minus Control Matching (Decreases) |
| 5070124 | Healthy Controls | 40 | WCST > Control |
| 5070125 | MRMD | 5 | WCST – WCS Control, Lupron |
| 5070125 | MRMD | 5 | WCST – WCS Control, Lupron and Progesterone |
| 5070125 | MRMD | 5 | WCST – WCS Control, Lupron and Estrogen |
| 5070137 | Healthy Controls | 12 | WCST - Control, Activations |
| 5070137 | Healthy Controls | 12 | WCST - Control, Deactivations |
| 5070141 | Healthy Controls | 21 | Negative - Neutral Feedback |
| 5070148 | Healthy Controls | 36 | Inhibition - Control, No Additional Task Knowledge |
| 5070148 | Healthy Controls | 16 | Inhibition - Control, Additional Task Knowledge |
| 5070156 | Healthy Controls | 9 | Matching After Negative Feedback - Control Matching |
| 5070156 | Healthy Controls | 9 | Matching After Positive Feedback - Control Matching |
| 5070156 | Parkinson | 8 | Matching After Negative Feedback - Control Matching |
| 5070156 | Parkinson | 8 | Matching After Positive Feedback - Control Matching |
| 5070158 | Healthy Controls | 18 | Matching WCST vs. Matching |
| 5070159 | Healthy Controls | 6 | Set Shifting Task |
| 5070160 | Healthy Controls | 6 | WCST > Matching, Young |
| 5070160 | Healthy Controls | 6 | WCST > Matching, Elderly |
| 5070170 | Schizophrenia | 15 | WCST - Rest |
| 5070170 | Healthy Controls | 15 | WCST – Rest |
| 5080215 | Healthy Controls | 10 | WCST |
| 7100280 | Healthy Controls | 12 | (Discrimination based on different dimension) - (Discrimination based on same dimension) |

| | | | |
|---|---|---|---|
| 8010020 | Healthy Controls | 12 | Negative Feedback |
| | | | Positive Feedback |
| 17030047 | Healthy Controls | 30 | Controls: Activated Brain Regions During first error feedback |
| | IBS | 30 | IBS: Activated Brain Regions During First error feedback |
| 17030049 | Healthy Controls | 19 | Matching after negative feedback > Matching after control feedback, p<.05 |
| | | | Matching after positive feedback > Matching after control feedback, p<.05 |
| 17030050 | Healthy Controls | 14 | Second Negative Feedback > Second Positive Feedback |
| | | | Second Negative Feedback < Second Positive Feedback |
| | SLE | 14 | Second Negative Feedback > Second Positive Feedback |
| | | | Second Negative Feedback < Second Positive Feedback |
| 17030051 | Healthy Controls | 15 | Set Shifting Error Feedback vs First Correct Feedback of 5 Consecutive Corrects |
| | AN | 15 | Set Shifting Error Feedback vs First Correct Feedback of 5 Consecutive Corrects |
| 17030053 | Healthy Controls | 16 | 2-Negative Feedback vs 1-Negative Feedback, p < 0.05 FDR corrected |
| 17040059 | Healthy Controls | 176 | 1st shift - 2nd shift during the WCST |
| 17040060 | Healthy Controls | 31 | Dimension change - Dimension repeat |
| 17040061 | Healthy Controls | 12 | (No instruction of dimensional change) – (Instruction of dimensional change) |
| 17040063 | Healthy Controls | 14 | WCST-Rest |

Cognitive term used was “Paradigm Class: Wisconsin Card Sorting Test (WCST)”.
Subjects included were healthy controls, Menstrually-related mood disorder (MRMD), Parkinson, Schizophrenia, Irritated bowel syndrome (IBS), systemic lupus erythematosus (SLE), and Anorexia Nervosa (AN).

**Table S4. List of experiments included for meta-analysis of activation coordinates reported for no-go continuous task performance.**

| BrainMap ID | Group Type | Sample Size | Experiment |
|---|---|---|---|
| 30226 | Healthy Controls | 15 | Generic Go/No-Go Activation |
| 30336 | Healthy Controls | 11 | Specific Activation Areas During NO-GO Phase |
| 30348 | Healthy Controls | 14 | Successful NoGos |
| 30351 | Healthy Controls | 9 | Go/NoGo Task - Control Task |
| 30354 | Healthy Controls | 16 | Correct NoGo – Go |
| 30356 | Healthy Controls | 14 | Go/NoGo – Go |
| 30362 | Healthy Controls | 19 | Go/NoGo - Go, Developing Controls |
| 5070119 | Healthy Controls | 17 | Response Inhibition |
| 5070122 | Healthy Controls | 42 | Response Inhibition |
| 5070134 | Healthy Controls | 21 | Activations For Correct Inhibitions |
| 5070136 | Healthy Controls | 14 | Response Inhibition |
| 5070139 | Healthy Controls | 15 | Cued vs. Uncued, Increases |
| | | | Cued vs. Uncued, Decreases |
| 5070140 | Healthy Controls | 21 | Go/No-Go > Go |
| 5070144 | Healthy Controls | 15 | Fast and Slow Successful Response Inhibitions |
| 5070147 | Healthy Controls | 5 | No-Go Dominant Foci |
| 5070153 | Healthy Controls | 6 | Go/No-Go vs. Go |
| 5070157 | Healthy Controls | 48 | Primary No-Go Effects |
| | Healthy Controls | 28 | Primary Counting No-Go Effects |
| 5070176 | Healthy Controls | 8 | Increases |
| | | | Decreases |
| | | | Linear Increases With Number of Trials Equated Per Block |
| | | | Linear Decreases With Number of Trials Equated Per Block |
| 5120247 | Healthy Controls | 16 | Event-Related STOPS |
| 5120248 | Healthy Controls | 6 | No-Go Dominant Area |
| 5120249 | Healthy Controls | 14 | Low Probability No-Gos, Correct Rejects |
| 7080203 | Healthy Controls | 14 | Response Inhibition |
| | OCD | 6 | Response Inhibition |
| 7080204 | Healthy Controls | 20 | NoGo, Successful Inhibition - NoGo, Unsuccessful Inhibition |

| | | | |
|---|---|---|---|
| 7080219 | OCD | 14 | Correct Inhibition |
| | Healthy Controls | 14 | Correct Inhibition, Normals |
| 7090271 | Healthy Controls | 23 | Go/NoGo Task, Adults |
| | Healthy Controls | 29 | Go/NoGo Task, Adolescents |
| 7110343 | Healthy Controls | 27 | Activity during Movement Preparation (NoGo) |
| 8080236 | Healthy Controls | 30 | No-Go Activations |
| 9020035 | PTSD | 23 | No Go - Go |
| | Healthy Controls | 23 | No Go - Go |
| | Trauma-exposed Patients | 17 | No Go - Go, |
| 9020051 | ADHD | 25 | No Go – Fixation |
| | Healthy Controls | 25 | No Go - Fixation |
| 9050074 | Bipolar Disorder | 11 | NoGo > Go |
| | Healthy Controls | 13 | NoGo > Go |
| 9100172 | Healthy Controls | 10 | No Go vs. Go, Maniac State |
| | | | No Go vs. Go, Healthy Controls, Euthymic State |
| | Bipolar DIsorder | 10 | No Go vs. Go, Bipolar Disorder Patients, Maniac State |
| | | | No Go vs. Go, Bipolar Disorder Patients, Euthymic State |
| 9100173 | Healthy Controls | 20 | NoGo > Go, Correct Responses |
| | Bipolar Disorder | 20 | NoGo > Go, Correct Responses |
| 10010001 | Healthy Controls | 22 | Disjunction Analysis, Stop vs. Uncertain-Go but not Uncertain-Go vs. Certain-Go |
| 10030047 | Healthy Controls | 16 | Go Trials vs. No-Go Trials |
| | Manic Patients | 16 | Go Trials vs. No-Go Trials |
| 10030071 | Bipolar DIsorder | 27 | Go/No-go > Fixation |
| | Healthy Controls | 28 | Go/No-go > Fixation |
| 10030074 | Healthy Controls | 18 | No-Go - Go (Go/No-Go) |
| 10030081 | Healthy Controls | 11 | NoGo vs. Fixation |
| | Parkinson’s Disease | 15 | NoGo vs. Fixation |
| 10030084 | Healthy Controls | 11 | No/Go - Go, Correct Trials |
| | PTE | 7 | No/Go - Go, Correct Trials |
| 10060114 | Healthy Controls | 17 | Activation in Response to Correct Rejections |
| | MDD | 16 | Activation in Response to Correct Rejections |
| 10080149 | Healthy Controls | 25 | No-Go vs. Frequent-Go |

| | | | |
|---|---|---|---|
| | | | No-Go vs. Infrequent-Go |
| 10080157 | Healthy Controls | 11 | No-Go > Go |
| | Unaffected Siblings | 11 | No-Go > Go |
| | ADHD | 11 | No-Go > Go |
| 10080161 | Healthy Controls | 13 | NoGo Inhibition vs. Go, Experimental Session |
| 10080190 | Healthy Controls | 11 | No-Go > Go |
| 10080193 | Healthy Controls | 12 | No Go vs. Go, Independently of MAO-A Genotype |
| 13020010 | Healthy Controls | 21 | Correct NoGo Trials vs. Correct Go Trials |
| | Schizophrenia | 21 | Correct NoGo Trials vs. Correct Go Trials |
| 13020014 | Pediatric Bipolar Disorder | 26 | NoGo - Go |
| | Healthy Controls | 22 | NoGo - Go |
| 13020016 | Healthy Controls | 30 | NoGo - Go |
| | Bipolar Disorder | 32 | NoGo - Go |
| 14050049 | Healthy Controls | 12 | Go/No-Go |
| | ASD | 10 | Go/No-Go |
| 14080159 | Healthy Controls | 12 | Regional activation for the NoGo vs Go trials, All subjects |
| | ASD | 12 | |
| 14120258 | PTSS | 16 | No-Go vs Go |
| | Healthy Controls | 14 | No-Go vs Go |
| 15100134 | Healthy Controls | 20 | Initial Go/No-go > Tonic Alertness |
| | | | Complex Go/No-go > Tonic Alertness |
| 15110175 | Healthy Controls | 102 | Nogo > Go |
| 16030070 | Healthy Controls | 13 | Correct NoGo Trials - Correct Go Trials in All Subjects |
| | ADHD | 16 | |
| 16030073 | ADHD | 15 | Go/NoGo Task |
| | Healthy Controls | 15 | Go/NoGo Task |
| 16040099 | Healthy Controls | 14 | Significant Activations During the Response Inhibition Task in All Subjects |
| | 22q11.2DS | 13 | |
| | IDD | 9 | |
| 16060140 | Healthy Controls | 10 | Pre vs. Post training, Cognitive Training Group, Response inhibition |
| 16120232 | Healthy Controls | 14 | No-Go > Oddball, Controls, Sham |
| | | | No-Go > Oddball, Controls, ATD |

| | | | |
|---|---|---|---|
| | ASD | 14 | No-Go > Oddball, ASD, Sham |
| | | | No-Go > Oddball, ASD, ATD |
| 16120234 | Healthy Controls | 15 | Letter No-Go - Letter Go, Controls |
| | ASD | 15 | Letter No-Go - Letter Go, Autistic Subjects |
| 17050107 | Healthy Controls | 14 | no-go: healthy controls |
| | ADHD | 17 | no-go: ADHD |
| 17080165 | Healthy Controls | 18 | No-go > Informatively cued (go) |
| 17080184 | Healthy Controls | 24 | go/no go task: no-go > go; control group |
| 17090198 | Healthy Controls | 12 | Typical, Isolated > Baseline |
| | Dyslexia | 12 | Dyslexia, Isolated > Baseline |
| 17100229 | Healthy Controls | 31 | lure > no go: controls |
| 18010011 | Healthy Controls & Pedophilia | 7/11 | No-Go > Go, All Participants |

Cognitive term used was “Paradigm Class: Go/No-Go”.
Subjects included were healthy controls, Obsessive-compulsive disorder (OCD), Post-traumatic stress disorder (PTSD), Trauma-exposed patients, attention-deficit hyperactivity disorder (ADHD) and unaffected siblings, Bipolar, Manic patients, Parkinson, prenatal tobacco exposed subjects, Schizophrenia, Major Depressive Disorder Patients, Autism Spectrum Disorder, Dyslexia, 22q11.2 Deletion Syndrome, Idiopathyic developmental Disability (IDD).

**Table S5. List of experiments included for meta-analysis of activation coordinates reported for fluid intelligence.**

| BrainMap ID | Group Type | Group Size | Cognitive Task | Experiment |
|---|---|---|---|---|
| 30202 | Healthy Controls | 8 | Conditional Reasoning | Posttest - Pretest |
| 30282 | Healthy Controls | 8 | Matrix Reasoning | Linear Trend Analysis for Relational Complexity |
| | | | | Linear Trend Analysis for Distractor |
| | | | | Complexity levels 3-4 Minus Distractor levels 3-4 |
| 6050065 | Healthy Controls | 7 | Pattern Matching | Rest vs. Tasks |
| | Healthy Controls | 8 | | |
| | Healthy Controls | 10 | | |
| | Healthy Controls | 7 | | |
| | Healthy Controls | 10 | | |
| 7070175 | Healthy Controls | 8 | Perceptual Maze | Perceptual Maze vs. Motor Control, Increase in rCBF |
| | | | | Perceptual Maze vs. Motor Control, Decrease in rCBF |
| 7110309 | Healthy Controls | 20 | Conditional Reasoning | Verification: 2-Mismatch > 1-Mismatch > 0-Mismatch |
| | | | | Falsification: 2-Mismatch > 1-Mismatch > 0-Mismatch |
| | | | | Affirmative Throughout: 1-Mismatch > 0-Mismatch |
| | | | | Verification, Hits Only: 2-Mismatch > 1-Mismatch > 0-Mismatch |
| 8040095 | Healthy Controls | 15 | Transitive Inference | (Novel Sequence Pairs > Previously Seen Sequence) > (Novel Nonoverlapping Pairs > Previously Seen Nonoverlapping Pairs), Brain Activations During Transitive Inference Condition |
| | Schizophrenia | 15 | Transitive Inference | (Novel Sequence Pairs > Previously Seen Sequence) > (Novel Nonoverlapping Pairs > Previously Seen Nonoverlapping Pairs), Brain Activations During Transitive Inference Condition |
| 12070050 | Healthy Controls | 15 | Transitive Inference | Transitive Inference - Visual Height Comparison |
| 13070060 | Healthy Controls, High IQ | 22 | Geometric Analogical Reasoning Task | Task Difficulty (Diagonal > Horizontal > Vertical > Identity), All subjects |
| | Healthy Controls, Low IQ | 18 | | |
| 14010003 | Healthy Controls | 12 | Pattern Matching | Induction Conjunction Analysis |

| | | | | |
|---|---|---|---|---|
| | | | | Visualization Conjunction Analysis |
| | Healthy Controls | 10 | Pattern Matching | Induction Conjunction Analysis |
| | | | | Visualization Conjunction Analysis |
| 14010009 | Healthy Controls | 16 | Non-verbal Reasoning | Peak Activation Coordinates During Reasoning |
| | | | | Main Effect of Rule Complexity for Simultaneous Panels |
| | | | | Main Effect of Analogical Distance for Simultaneous Panels |
| | Healthy Controls | 21 | Non-verbal Reasoning | Main Effect of Rule Complexity for Separate Panels |
| | | | | Main Effect of Analogical Distance for Separate Panels |
| 14010023 | Healthy Controls | 8 / 10 | Matrix Reasoning | Activations, All Subjects |
| | Healthy Controls | 10 | | |
| 14030041 | Healthy Controls | 14 | Non-verbal reasoning | Areas modulated by task difficulty (Hard > Easy) |
| | | | | Areas modulated by response correctness (Correct > Incorrect) |
| 14050115 | ASD | 12 | Embedded Figures Task | Embedded Figures Task > Control task |
| | Healthy Controls | 12 | Embedded Figures Task | Embedded Figures Task > Control Task |
| 14050128 | Healthy Controls | 15 | Pattern Matching | Pattern Matching > Fixation, Non-Autistic Controls |
| | | | Matrix Reasoning | Raven’s Standard Progressive Matrices > Fixation |
| | | | | Fixation > Raven’s Standard Progressive Matrices |
| | ASD | 18 | Pattern Matching | Pattern Matching > Fixation |
| | | | | Fixation > Pattern Matching |
| | | | Matrix Reasoning | Raven’s Standard Progressive Matrices > Fixation |
| | | | | Fixation > Raven’s Standard Progressive Matrices |
| 14090184 | Healthy Controls | 26 | Numerosity discrimination task | Main Effect of Numerosity Task |
| 14090185 | Healthy Controls | 20 | Non-symbolic reasoning | Nonsymbolic > Symbolic |
| | | | | Dot-dot > cross-notation trials |
| 14090187 | Healthy Controls | 19 | Non-symbolic reasoning | (nonsymbolic - control) - (symbolic - control) |

| | | | | |
|---|---|---|---|---|
| 14090190 | Healthy Controls | 33 | Non-symbolic reasoning | Nonsymbolic: Numbers in numerical order > Numbers sorted by luminance |
| | | | | (Dots in numerical order > Dots sorted by luminance) AND (Dot Array with largest numerosity > Brightest Dot Array) |

Cognitive term used was Problem Solving.

In total, 10 different types of task were selected: Conditional Reasoning, Matrix Reasoning, Pattern Matching, Perceptual Maze, Transitive Inference, Geometric Analogical Reasoning Task, Non-verbal Reasoning, Embedded Figures Task, Numerosity discrimination task, Non-symbolic reasoning.

All these tasks have in common the fact that they involve problem solving without the need of previous knowledge acquisition.

Subjects included were healthy controls, with different IQ levels Schizophrenia, Autism Spectrum Disorder (ASD).

**Table S6. List of experiments included for meta-analysis of activation coordinates reported for reading decoding.**

| BrainMap ID | Group Type | Sample Size | Experiment |
|---|---|---|---|
| 30027 | Healthy Controls | 13 | Encoding vs. Reference, Increases |
| | | | Encoding vs. Reference, Decreases |
| 30081 | Stuttering | 10 | Solo vs. Rest, Activations |
| | | | Solo vs. Rest, Deactivations |
| | Healthy Controls | 10 | Solo vs. Rest, Activations |
| | | | Solo vs. Rest, Deactivations |
| 30216 | Healthy Controls | 10 | Regular Characters vs. Fixation |
| | | | Irregular Characters vs. Fixation |
| 30241 | Healthy Controls | 16 | Read Words Aloud - Words Control |
| | | | Words Control - Read Words Aloud |
| 30255 | Healthy Controls | 11 | Word Reading - Fixation |
| | | | Fixation - Word Reading |
| 30257 | Healthy Controls | 11 | Aloud - Silent Words |
| | | | Silent - Aloud Words |
| 30258 | Healthy Controls | 10 | Irregular + Regular - Zero-Speak |
| 30260 | Healthy Controls | 8 | Word Identification > Fixation |
| | | | Fixation > Word Identification |
| 30265 | Healthy Controls | 14 | Irregular Pronunciation - Fixation |
| | | | Fixation - Irregular Pronunciation |
| 30317 | Stuttering | 4 | Overt Solo – Rest |
| | | | Rest - Overt Solo |
| | Healthy Controls | 4 | Overt Solo – Rest |
| | | | Rest - Overt Solo |
| 30378 | Healthy Controls | 10 | Oral Reading - Baseline, Controls |

| | | | |
|---|---|---|---|
| | Stuttering | 13 | Oral Reading - Baseline, Pre-Treatment |
| | | | Oral Reading - Baseline, Post-Treatment |
| | | | Oral Reading - Baseline, 1 Year |
| 4020008 | Healthy Controls | 12 | Word Reading vs. See and Say, Increases |
| | | | Word Reading vs. See and Say, Decreases |
| 4020011 | Healthy Controls | 6 | Reading - Feature, Listening to words at 150 + 1000 ms duration |
| 4020012 | Healthy Controls | 6 | Reading words aloud at 40 wpm - Listening to words at 40 wpm |
| 4020013 | Healthy Controls | 6 | Aloud - Rest |
| 5070163 | Healthy Controls (Language I) | 6 | Main Effect of Reading |
| | Healthy Controls (Language II) | 6 | |
| 5070171 | Healthy Controls | 15 | Kana and Kanji Conjunction |
| 6050051 | Healthy Controls | 10 | Encoding, High Imagery, Foreign Language vs. Reference |
| | | | Encoding, High Imagery, Native Language vs. Reference |
| | | | Encoding, Low Imagery, Foreign Language vs. Reference |
| | | | Encoding, Low Imagery, Native Language vs. Reference |
| 7020036 | Healthy Controls | 12 | Word Reading vs. Fixation |
| 7110316 | Healthy Controls | 36 | Read > False Fonts |
| 8110263 | Healthy Controls | 10 | Bisyllables > Monosyllables |
| 9010023 | Healthy Controls | 9 | High Frequency Regular Words vs. Rest |
| | | | Low Frequency Regular Words vs. Rest |
| | | | High Frequency Exception Words vs. Rest |
| | | | Low Frequency Exception Words vs. Rest |
| | Semantic Dementia | 5 | High Frequency Regular Words vs. Rest |
| | | | Low Frequency Regular Words vs. Rest |
| | | | High Frequency Exception Words vs. Rest |
| | | | Low Frequency Exception Words vs. Rest |

| | | | |
|---|---|---|---|
| 9030063 | Healthy Controls | 14 | Brain Activation During Read Task |
| | | | Brain Deactivation During Read Task |
| 9030066 | Healthy Controls | 32 | Locations of Significant Maxima for fMRI Study |
| 9090121 | Healthy Controls | 14 | Reading Aloud > Word Recognition, Kanji |
| | | | Reading Aloud > Word Recognition, Kana |
| 11010016 | Parkinson | 10 | Paragraph Reading > Rest, Pre-Lee Silverman Voice Treatment LOUD |
| | | | Paragraph Reading > Rest, Post-Lee Silverman Voice Treatment LOUD |
| 16030053 | Healthy Controls | 16 | Reading Aloud-All words, LFS > HFS |
| 16030072 | Healthy Controls | 16 | All Conditions vs. Fixation |

Cognitive term used was “Paradigm Class: Reading (Overt)”.
Subjects were instructed to read overtly words (individual, paragraphs, disyllables, characters), Subjects included were healthy controls, or suffered stuttering (at different treatment stages), semantic dementia and Parkinson’s disease.

**Table S7. List of experiments included for meta-analysis of activation coordinates reported for vocabulary comprehension task performance.**

| BrainMap ID | Group Type | Group Size | Experiment |
|---|---|---|---|
| 30005 | Healthy Controls | 13 | Unprimed Semantic Category Decision - Baseline |
| 30006 | Healthy Controls | 12 | Words > Tones - Active |
| 30018 | Healthy Controls | 11 | Semantic > Episodic |
| 30019 | Healthy Controls | 11 | Semantic - Episodic |
| 30079 | Healthy Controls | 13 | Visual Meaning - Control |
| | | | Auditory Meaning - Control |
| 30080 | Healthy Controls | 37 | Same vs. Different Word, Independent of Case |
| 30091 | Healthy Controls | 5 | Body Parts > Numbers (Block Design) |
| | Healthy Controls | 6 | Body Parts > Numbers (Event-Related Design) |
| 30136 | Healthy Controls | 12 | Semantic > Nonsemantic and Intentional Learning |
| 30141 | Healthy Controls | 12 | Semantic Processing - Mouse-Click |
| 30164 | Healthy Controls | 8 | Semantic > Phonological (Double Subtraction) |
| 30167 | Healthy Controls | 20 | Synonym AND Easy Categorization |
| | | | Synonym AND Hard Categorization |
| 30189 | Healthy Controls | 6 | Category Only Subjects: Categorization - Control |
| | Healthy Controls | 6 | Mixed Subjects: Categorization - Control |
| 30213 | Healthy Controls | 6 | Semantic Relatedness Judgment vs. Crosshair Fixation |
| 30232 | Healthy Controls | 12 | Activation Specific to Semantic Decision |
| 30234 | Healthy Controls | 12 | Specific to Retrieving Knowledge about Tools |
| 30334 | Healthy Controls | 10 | Release Semantic |
| | | | Hold Semantic |
| 30347 | Healthy Controls | 12 | Semantic > Phonological |
| 30350 | Healthy Controls | 20 | Semantic - Lexical |
| 30365 | Healthy Controls | 20 | Nonsocial Semantic vs. Control |
| 30424 | Healthy Controls | 17 | Correct vs. Incorrect Source Memory During Encoding |
| 30426 | Healthy Controls | 10 | English Words - Control, SGP |
| | | | Mandarin Characters - Control, SGP |
| | Healthy Controls | 9 | English Words - Control, PRC |
| | | | Mandarin Characters - Control, PRC |

| | | | |
|---|---|---|---|
| 30431 | Bipolar | 5 | Semantic Decision > Fixation |
| | | | Fixation > Semantic Decision |
| 30432 | Healthy Controls | 5 | Semantic Decision – Control |
| | Schizophrenia | 5 | Semantic Decision – Control |
| 30434 | Healthy Controls | 9 | Words – Phonemes |
| 30461 | Healthy Controls | 6 | Activation Areas |
| 5040019 | Healthy Controls | 10 | Semantic Judgment - Anticipation (Activations) |
| | | | Semantic Judgment - Anticipation (Deactivations) |
| 5040026 | Healthy Controls | 6 | Words: Associative and Visual Semantics vs. Baseline |
| | | | Pictures: Associative and Visual Semantics vs. Baseline |
| 5040055 | Healthy Controls | 15 | Abstract Concepts > Sounds, Visual Attributes, and Hand Movements |
| 5040061 | Healthy Controls | 12 | Chinese – Baseline |
| | | | English – Baseline |
| 5070094 | Healthy Controls | 6 | Natural Black and White Relative to Man-Made Black and White (Matching) |
| 5080182 | Healthy Controls | 24 | Semantic Decision on Words |
| 5080190 | Healthy Controls | 6 | Semantic – Phonological |
| 5080191 | Healthy Controls | 8 | Objects Minus Letter-Spatial |
| 6050053 | Healthy Controls | 12 | Associative Word Learning vs. Shallow Single Word Encoding |
| 6060077 | Healthy Controls | 26 | Word > Face |
| | Schizophrenia | 17 | Word > Face |
| 7020040 | Healthy Controls | 6 | Noun-Noun Comparisons vs. Rest |
| | | | Verb-Noun Comparisons vs. Rest |
| 7020055 | Healthy Controls | 12 | Object Identity > Plus-Minus |
| 7020066 | Healthy Controls | 9 | Verb-Noun Comparison vs. Rest |
| | | | Rest vs. Verb-Noun Comparison |
| 7050138 | Healthy Controls | 11 | Label Pictures > Match Pictures |
| 7070170 | Healthy Controls | 12 | Language vs. Tones |
| 7070188 | Healthy Controls | 50 | Semantic Categorization vs. Perceptual Categorization |
| 7090260 | Healthy Controls | 6 | Chinese Semantic vs. Asterisks |
| | | | English Semantic vs. Asterisks |
| 7100276 | Healthy Controls | 16 | Simple – Fixation |
| 7110336 | Healthy Controls | 14 | Picture > Fixation |

| | | | |
|---|---|---|---|
| | | | Chinese Character > Fixation |
| 7120358 | Healthy Controls | 20 | Abstract/Concrete > Good/Bad |
| 7120385 | Healthy Controls | 15 | (Coherent Explicit + Coherent Implicit + Incoherent) > Pseudowords |
| 8040092 | Healthy Controls | 16 | Recall > No Recall |
| | Schizophrenia | 16 | Recall > No Recall |
| 8040099 | Healthy Controls | 18 | Correct Associated |
| | | | Correct Non Associated |
| 8050117 | Healthy Controls | 21 | Deep vs. Shallow Encoding |
| | Siblings of Schizophrenia Patients | 38 | Deep vs. Shallow Encoding |
| 8050132 | Healthy Controls | 9 | Deep vs. Shallow Encoding |
| 8060151 | Healthy Controls | 17 | Regions Active at the Target Word During Relational Trials Relative to Item Specific Trials |
| 8080205 | Healthy Controls | 12 | Successful Encoding > Baseline Triad |
| 8080229 | Healthy Controls | 14 | Picture, Conception > Rest |
| 9090125 | Healthy Controls | 12 | Naming > Letter Strings |
| | ASD | 12 | Naming > Letter Strings |
| 9090154 | Healthy Controls | 14 | SYN > STRINGS |
| | Primary Progressive Aphasia | 14 | SYN > STRINGS |
| 10080210 | Healthy Controls | 14 | 4 Target > 2 Target |
| | | | Weak > Strong, Associative Strength |
| 11010025 | Healthy Controls | 12 | Overall Semantic Contrast |
| 11010031 | Healthy Controls | 14 | [Semantic-Lexical] & [Semantic-Phonological] Conjunction Analysis, Activations |
| 11080061 | Healthy Controls | 12 | All Words vs. Letter Strings |
| 11110113 | Healthy Controls | 12 | Tools vs. Letter Strings |
| | | | Animals vs. Letter Strings |
| | | | Tool Actions vs. Letter Strings |
| | | | Biological Actions vs. Letter Strings |
| 12020002 | Healthy Controls | 12 | Low > High Frequency, Block Design, Size |
| | Healthy Controls | 8 | Low > High Frequency, Block Design, Fixation |
| | Healthy Controls | 12 | Low > High Frequency, Event-Related Design |
| 12070055 | Healthy Controls | 16 | Running Verbs > Wingdings |

| | | | |
|---|---|---|---|
| | | | Speaking Verbs > Wingdings |
| | | | Hitting Verbs > Wingdings |
| | | | Cutting Verbs > Wingdings |
| | | | Change of State Verbs > Wingdings |
| 12070059 | Healthy Controls | 12 | Words (Nouns + Verbs) - Letter Strings |
| 12070061 | Healthy Controls | 20 | (Nouns - Fixation) > (Verbs - Fixation) |
| | | | (Verbs - Fixation) > (Nouns - Fixation) |
| 12100071 | Healthy Controls | 10 | Semantic Jugdment > Fixation |
| 13030019 | Healthy Controls | 13 | Words (Nouns + Verbs) > Non-words |
| 13030030 | Healthy Controls | 12 | Literal > Rest |
| 14010002 | Healthy Controls | 19 | Categorical Knowledge (Non-Social > Social) |
| 16030050 | Healthy Controls | 13 | Meaning - Lines |
| | Healthy Controls | 8 | Meaning - Lines |
| 16030057 | Healthy Controls | 8 | Abstract/Concrete > Syllable |
| | | | Abstract/Concrete > Case |
| 16030062 | Healthy Controls | 35 | Related Words > Baseline |
| | | | Unrelated Words > Baseline |
| 16030067 | Healthy Controls | 12 | High Confidence Hits > Misses, Explicit Learning |
| | Healthy Controls | 12 | High Confidence Hits > Misses, Explicit Learning |
| 16060153 | Healthy Controls | 15 | Social + Animal > Numbers |
| 17010014 | Healthy Controls | 16 | Semantic > Rhyming, All subjects |
| | Dyslexia | 16 | |
| 17060113 | Healthy Controls | 10 | Semantic word matching |
| | Healthy Controls | 14 | Semantic word matching |
| | Dyslexia | 11 | Semantic word matching |
| | Dyslexia | 11 | Semantic word matching |
| 17090204 | Healthy Controls | 14 | Incongruent > Congruent |
| 60100159 | Healthy Controls | 14 | Deep Word Encoding - Shallow Word Encoding |
| | Schizophrenia | 14 | Deep Word Encoding - Shallow Word Encoding |
| 60100164 | Healthy Controls | 22 | Semantic Self-Paced vs. Perceptual Self-Paced |
| | | | Semantic Fixed-Paced vs. Perceptual Fixed-Paced |

Cognitive term used was “Behavioral Domain is Cognition: Language Semantics”.
Subjects were instructed to a semantic decision (categorization/matching) based on input given in form of pictures or displayed/spoken words.
Subjects included were healthy controls, or suffered stuttering (at different treatment stages), semantic dementia and Parkinson’s disease.

**Movie S1. Brain volume animation of probabilistic activation maps estimated for card sorting and fluid intelligence, in addition to their respective activation overlap. Brain activation clusters representative of card sorting and fluid intelligence are colored in blue and green, respectively, whereas their overlapping regions are highlighted in red.**

**Movie S2. Brain volume animation of probabilistic activation maps estimated for reading decoding and fluid intelligence in addition to their respective activation overlap. Brain activation clusters representative of reading decoding and fluid intelligence are colored in yellow and green, respectively, whereas their overlapping regions are highlighted in red.**

**Movie S3. Brain volume animation of probabilistic activation maps estimated for reading decoding and no-go continuous performance, in addition to their respective activation overlap. Brain activation clusters representative of reading decoding and no-go continuous performance are colored in yellow and blue, respectively, whereas their overlapping regions are highlighted in red.**

**Movie S4. Brain volume animation of probabilistic activation maps estimated for reading decoding and vocabulary comprehension, in addition to their respective activation overlap. Brain activation clusters representative of reading decoding and vocabulary comprehension are colored in yellow and blue, respectively, whereas their overlapping regions are highlighted in red.**
